# Rat hepatitis E virus and novel paramyxoviruses in synanthropic rodents and shrews in Kenya

**DOI:** 10.64898/2026.04.21.719784

**Authors:** Griphin Ochola, Emilia Pulkkinen, Joseph G. Ogola, Henna Mäkelä, Moses Masika, Hanna Vauhkonen, Teemu Smura, Anne J. Jääskeläinen, Omu Anzala, Olli Vapalahti, Alois Wambua Mweu, Kristian M Forbes, Johanna F. Lindahl, Juha Laakkonen, Joni Uusitalo, Eda Altan, Essi M. Korhonen, Tarja Sironen

**Affiliations:** Department of Virology, Medicum, University of Helsinki, Helsinki, Finland; Virology and Immunology, HUS Diagnostic Center, University of Helsinki and Helsinki University Hospital, Helsinki, Finland; ISLAB Laboratory Centre, Kuopio, Finland; Department of Medical Microbiology and KAVI Institute of Clinical Research, University of Nairobi, Nairobi, Kenya; Department of Veterinary Biosciences, University of Helsinki, Helsinki, Finland; Department of Biological Sciences, University of Arkansas, Fayetteville, Arkansas, USA; Department of Medical Biochemistry and Microbiology, Uppsala University, Uppsala, Sweden; Lammi Biological Station, Faculty of Biological and Environmental Sciences, University of Helsinki, Lammi, Finland

**Author notes:** Equal contribution (Griphin Ochola, Emilia Pulkkinen).

**Keywords:** Zoonotic viruses, Rocahepevirus, jeilongvirus, NGS, *hepeviridae*, *Paramyxoviridae*

## Abstract

The majority of emerging infectious diseases are zoonotic, having their origin in wildlife before spilling over into the human population. While small mammals are recognized as critical reservoirs for these viruses, their viral diversity remains largely uncharacterized across many African countries. We conducted molecular surveillance of synanthropic rodents and shrews in the Kibera informal settlement in Nairobi and the rural Taita Hills region of Kenya to detect and characterize potential zoonotic viruses. Tissue samples from 228 rodents and shrews were screened for six viral families using PCR assays. Rat hepatitis E virus (HEV) (*Rocahepevirus ratti*), a rodent-associated virus with potential for human spillover, was identified in *Mus musculus* and *Rattus norvegicus* from Kibera. NGS was conducted for the HEV positive samples, and we obtained two near-complete HEV genomes from *Rattus norvegicus*, which clustered within rodent-associated HEV genotypes in the phylogenetic analysis. The two sequences from the *Rattus norvegicus* cluster together, indicating a close genetic relationship. Paramyxoviruses belonging to the genera *Jeilongvirus* and *Parahenipavirus* were detected both from Taita and Kibera in nine different samples from *Rattus norvegicus, Mus minutoides, Crocidura sp* and *Acomys ignitus*. One paramyxovirus positive sample (*Acomys ignitus*) from Taita was selected for further sequencing with NGS, and a complete genome of a new jeilongvirus was assembled. Phylogenetic analysis of the detected viruses confirmed the close relation to previously known rodent-borne jeilongviruses but also revealed potentially novel jeilong- and parahenipavirus species. Our findings highlight the circulation of potentially zoonotic viruses in both urban and rural small mammals in Kenya. It emphasizes the necessity of continued genomic surveillance of zoonotic viruses to mitigate risks of their spillover into human populations.

**Highlights:**

- Surveillance reveals diverse rodent-borne viruses circulating in Kenya.
- Rat-HEV was detected in *Rattus norvegicus* and *Mus musculus* from an urban low-income area.
- Paramyxoviruses were detected across multiple rodent and shrew species, including novel *Acomys ignitus* jeilongvirus.

**Graphical abstract:** 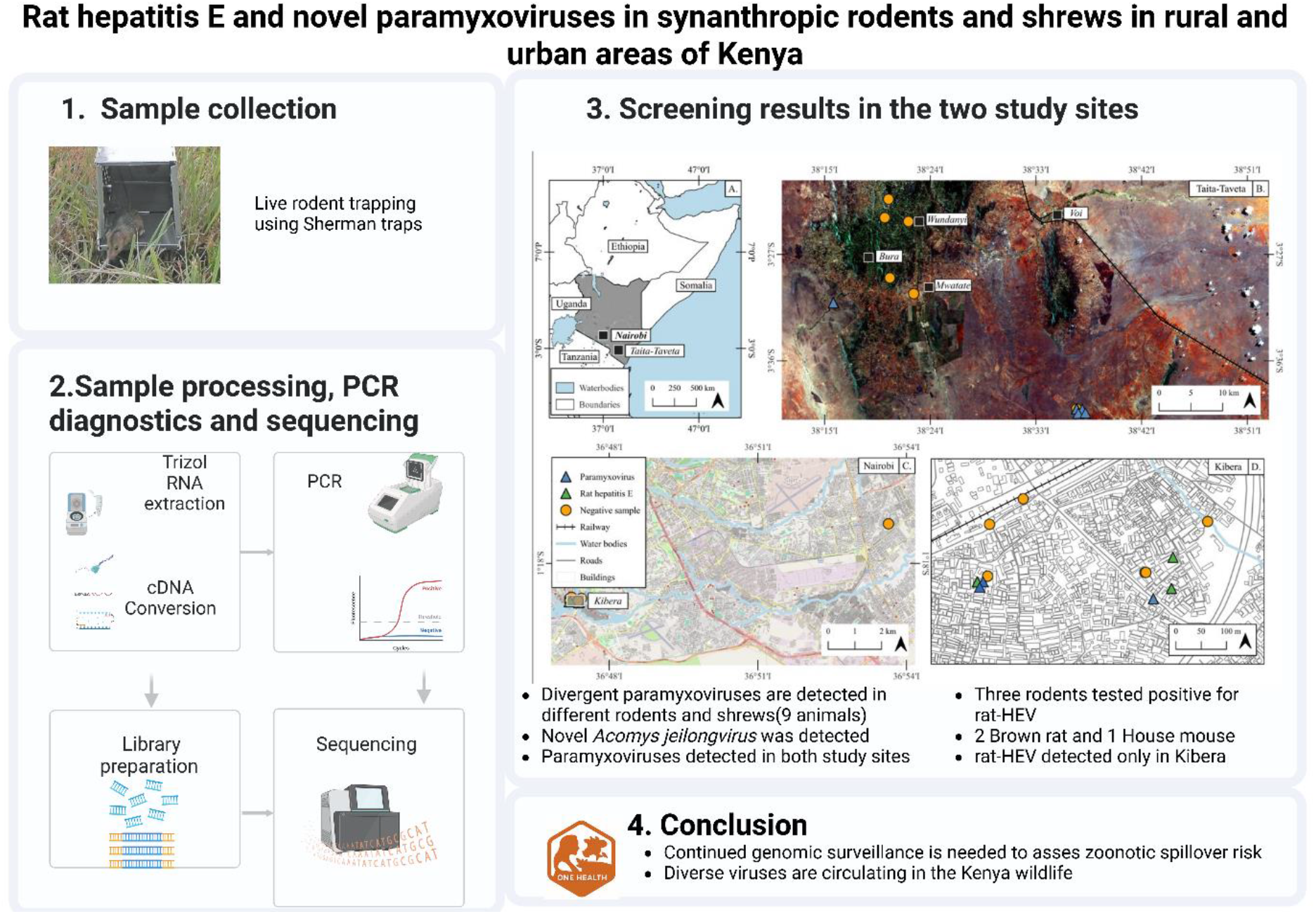

## Introduction

Emerging and re-emerging infections have become increasingly common in recent decades [1,2]. Their impact has affected public health, economies, and daily life, as seen clearly during the COVID-19 pandemic [3]. Most emerging infectious diseases (>60%) are zoonotic, and more than 70% of them originate from wildlife reservoirs. This highlights the critical role of wildlife in the maintenance and transmission of pathogens with epidemic and pandemic potential. Increasing globalization, environmental change, and intensified human–animal interactions further amplify the risk of pathogen emergence and spread[1,4]

Within the diverse spectrum of wildlife hosts, small mammals—most notably rodents and bats—function as primary reservoirs for a significant proportion of known zoonotic viral pathogens [5] These animals serve as natural reservoirs for several clinically relevant viral families, including *Hantaviridae, Arenaviridae*, and *Paramyxoviridae* families, as well as various orthopoxviruses [6–9]. Rodents’ ability to thrive in diverse environments, reproduce rapidly, and maintain long-term viral infections without exhibiting symptoms facilitate pathogen persistence and transmission [10,11]. Frequent interactions between small mammals, humans, and livestock further create continuous opportunities for viral spillover and zoonotic outbreaks.

Kenya’s geographical position and ecological characteristics creates an ideal hotspot for viral spillover, attributed to the high population of wildlife supporting an equally high diversity of zoonotic pathogens. Human activities and land use changes have been recorded as major triggers of viral spillovers from wildlife to human population [12]An increase in the population size of Kenya has led to the clearance of wildlife habitats for settlement and agriculture, increasing contact between wildlife and human [13].Currently, few studies have investigated rodent-borne pathogens in Kenya, but these have recorded both novel viruses and viruses with zoonotic potential: Kitale arenavirus, Lemniscomys striatus polyomavirus, Rat hepatitis E virus (rat-HEV) and paramyxoviruses [14–17]. A study of human samples from Kibera informal settlement demonstrated evidence of exposure of the locals to orthopox-, arena-and hantaviruses .

The lack of comprehensive studies on rodent-borne viruses in Kenya and East Africa, particularly across rural and urban settings, highlights the need for further research to better assess risks to human health. To fill this gap, this study was designed to conduct molecular surveillance of synanthropic rodents and shrews from Kibera (urban informal settlement) and Taita Hills (forest–agricultural interface) to (i) screen for selected viral families with zoonotic relevance, (ii) characterize detected virus genomes through sequencing and phylogenetic analyses, and (iii) evaluate their potential public health significance within a One Health framework.

## Material and Methods

### Study sites

The samples for this study were collected in 2016, 2017 and 2019 from broad locations; the Taita Hills region in Taita-Taveta county where sampling was conducted in April, August and October 2016, and May 2017, and Kibera informal settlement in Nairobi County, where sampling was conducted in February 2019. The Taita hills are heavily fragmented landscapes in the Eastern Arc mountains that are highly biodiverse and experiencing rapid land-use changes due to the high human population density around the hills [19]. The area has more natural environments and a significantly lower human population density compared to Kibera informal settlement, which is characterized by poor sanitation, limited infrastructure, and dense human population [20]. The exact location of the sites is shown on (Fig.1A). Detailed description of these two study sites is provided in the supplementary Material S1.

**Figure 1.**
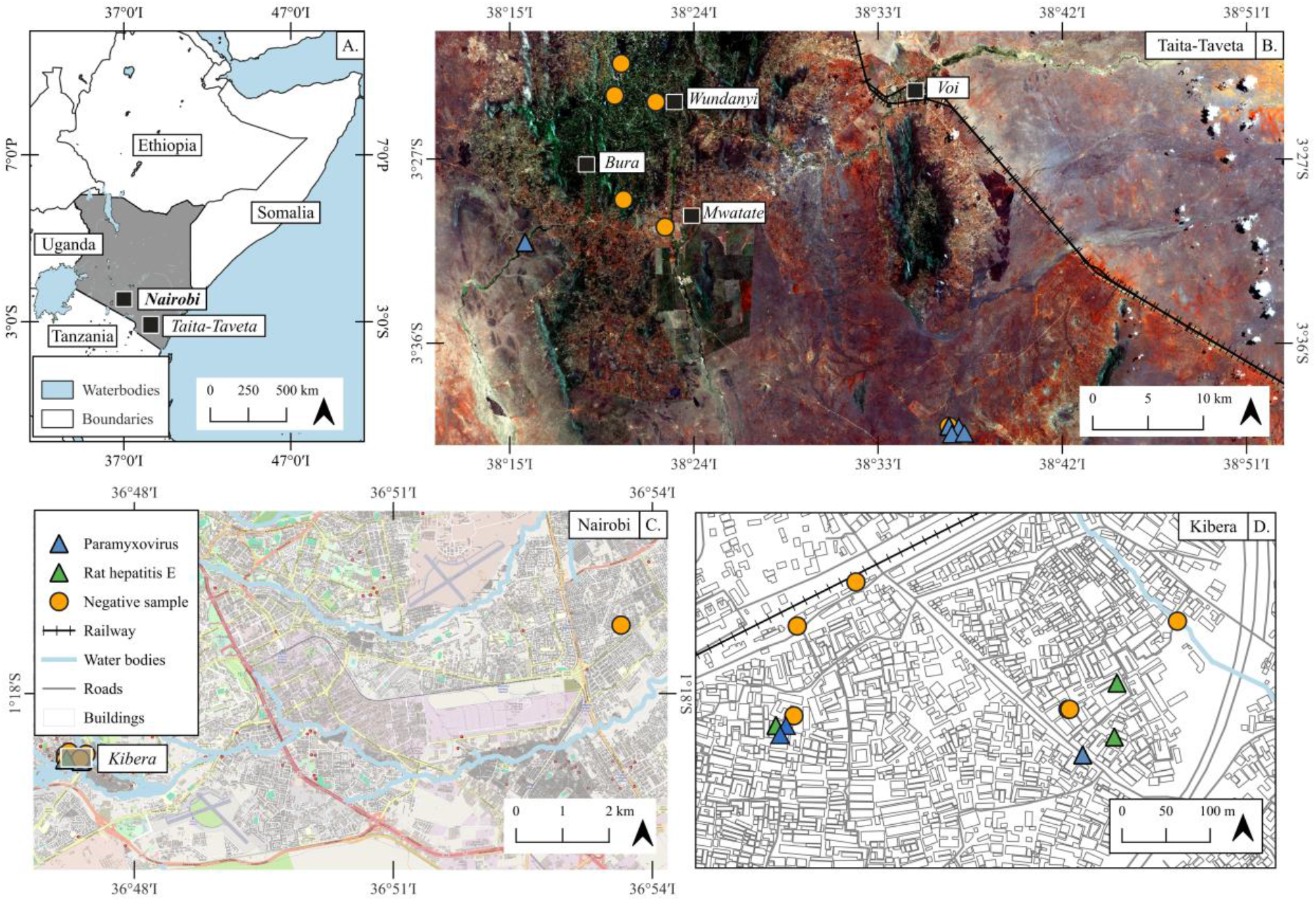
Maps of Kenya (A.), as a locator map and Taita Taveta (B.), Nairobi (C.) and Kibera (D.) showing the spatial location of sample sites where 228 samples were collected in 2016 and 2019. The blue and green triangles indicate paramyxovirus and HEV positive sites respectively, while orange dots indicate the negative sampling sites. The map was created using QGIS (version 3.36.2).

### Small mammal trapping and sample collection

In the Taita Hills, small mammals were captured using a stratified cross-sectional design. Line transects (∼100 m) were established in the homesteads and adjacent agricultural habitats, with 20 Sherman live traps placed at 5 m intervals and baited with oatmeal mixed with peanut butter. Traps were set for three nights at each point; they were set in the evening and checked at dawn; traps were closed during the day to prevent the animals entering and opened in the evening and re-baited.

Captured animals were transported to the Taita Research Station of University of Helsinki, anesthetized with an overdose of inhalation isoflurane, followed by cervical dislocation to ensure death. Individuals were identified morphologically to at least genus level, and sex, reproductive status, age class, body weight, head–body length, and tail length were recorded. Sampling in Kibera was done in the houses and around communal waste dumping sites using Sherman traps baited with dry fish, the rodents were dissected at University of Nairobi in the veterinary laboratory as described previously [18].

Tissues (lung, liver, kidney, spleen, intestine, and heart) were collected aseptically. Samples were preserved in RNAlater™ (Thermo Fisher Scientific, Waltham, MA, USA), except heart tissue, which was stored in phosphate-buffered saline (PBS; Gibco, Thermo Fisher Scientific). Samples were stored at −20°C during fieldwork, transferred to −80 °C at the University of Nairobi, and later shipped on dry ice to the University of Helsinki for molecular analyses.

All procedures were conducted in accordance with relevant national and institutional ethical guidelines; Biosafety, Animal Use and Ethics Committee of the University of Nairobi (permit No.FVM BAUEC/2018/180), the National Commission for Science, Technology and Innovation (permit No. NACOSTI/P/18/76501/22243) and the Kenya Wildlife Service (permit no. KWS/BRM/500). This study was approved by the University of Arkansas IACUC committee.

### PCR diagnostics

RNA was extracted from lung, liver, intestine, and kidney tissues using the TRIzol™ reagent (Thermo Fisher Scientific, Waltham, MA, USA) according to the manufacturer’s instructions and samples were stored at −80 °C. First-strand cDNA was synthesized from 5 µL of extracted RNA using the RevertAid First Strand cDNA Synthesis Kit (Thermo Fisher Scientific, Waltham, MA, USA).

Pathogen screening was performed using either RNA or cDNA templates with pathogen-specific nested, hemi-nested, RT-PCR, real-time PCR, or RT-qPCR assays as previously described for coronaviruses [21], paramyxoviruses [22], poxviruses [23], hantaviruses [24] and hepatitis E virus (HEV) [25], with minor modification for conventional PCR (supplementary material table 2). To acquire longer PCR products for Sanger sequencing, an additional conventional PCR was conducted for the paramyxovirus [26] and HEV [27] positive samples. Screening was conducted on selected organs based on pathogen tropism: lungs were tested for coronaviruses, paramyxoviruses, poxviruses, and hantaviruses; kidneys for paramyxoviruses; intestine for coronavirus; and liver for HEV.

PCR products of expected size were purified using the GeneJET Gel Extraction Kit (Thermo Fisher Scientific, Waltham, MA, USA) and subjected to Sanger sequencing. Obtained sequences were confirmed by BLAST analysis. Host species identity of positive animals was verified by cytochrome b gene sequencing of the cDNA as previously described[28].

### Sequencing and genome analysis

Samples for metagenomic NGS were selected based on the pathogen screening results. The NGS libraries were prepared using NEBNext® Ultra™ II RNA Library Prep Kit for Illumina® (New England Biolabs, Ipswich, MA, USA) system with slight modifications (full description in the Supplementary Material S2). NGS was performed at the Institute for Molecular Medicine Finland FIMM Genomics unit supported by HiLIFE and Biocenter Finland.

Raw reads were processed using the LazyPipe v3.1 metagenomic pipeline [29]. Quality filtering and human and mouse read removal were performed with fastp v.1.0.1 and BWA-MEM v.1.2.8, respectively[30,31], followed by de novo assembly using MEGAHIT [32]and homology searches were conducted with SANSparallel[33] UniprotKB database[34]. If de novo assembly resulted in fragmentary genomes, reads were mapped to reference viral genomes, and targeted PCR primers were designed to close gaps in the assemblies. Primer sequences and conditions are provided in Supplementary material s3.

Sequences in this study were aligned using Geneious Prime (Biomatters Ltd., Auckland, New Zealand), and MAFFT [35]. Maximum-likelihood phylogenetic trees were generated with IQ-TREE, using ModelFinder to determine the best-fitting substitution model prior to tree inference [36], and built with IQTREE [37] with 1000 ultrafast bootstrap replicates. Visualization was conducted using iTOL[38]. Open reading frames were acquired with Open Reading Frame Finder (NCBI).

## Results

A total of 228 small mammals of 11 species (nine rodent species and two shrew species) were collected in 2016,2017 and 2019 from the two sampling locations in Kenya as shown in (Fig.1). The distribution of the animals per site, Kibera informal settlement (n=195) and Taita hills area (n=33), is shown in Table 1.

**Table 1.**
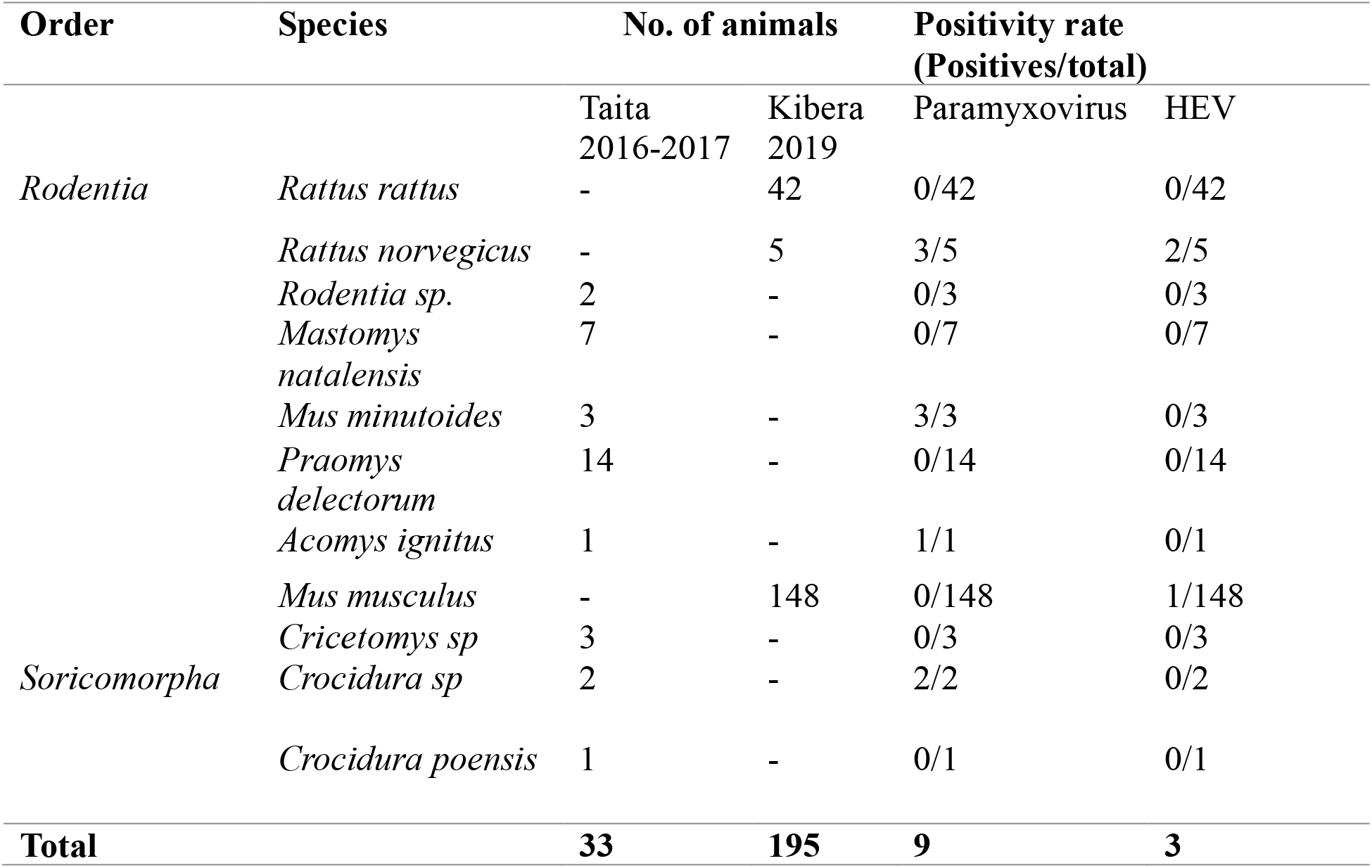
Numbers of individual animals captured from the two sampling sites, and the positivity rate of HEV and paramyxovirus per species.

Screening for the five pathogen groups identified viruses from two families: *Hepeviridae* and *Paramyxoviridae*. None of the samples were positive for coronaviruses, orthopoxviruses, or hantaviruses. Rat-HEV was detected exclusively in Kibera with a prevalence of 1.5% (3/195; 95% CI: 0.3–4.4%), while paramyxoviruses were detected in both study sites with a positivity rate of 1.5% (3/195; 95% CI: 0.3–4.4%) in Kibera and 18.1% (6/33; 95% CI: 7.0–35.5%) in Taita.

### Rat hepatitis E virus detection in the Kibera Informal settlement

Rat-HEV RNA was detected in liver tissues from three animals sampled in Kibera informal settlement: two *Rattus norvegicus;* KI49 and KI64 and one *Mus musculus*; KI137. The sequences obtained were submitted to GenBank (accession numbers: Supplementary Table 3). Partial RNA-dependent RNA polymerase sequences obtained by Sanger sequencing from the three rat-HEV–positive samples shared 99% nucleotide identity with each other. BLAST analysis showed 82–91% nucleotide identity to *Rocahepevirus ratti* sequences, with the highest identity (90–91%) to a strain reported from Guinea (GenBank accession PX408743).

Metagenomic sequencing yielded near-complete rat-HEV genomes from *Rattus norvegicus*, whereas no viral reads were recovered from the *Mus musculus* KI137. Comparative whole-genome analysis indicated that the two *Rattus norvegicus*-derived sequences were highly similar across the coding regions examined, sharing 99.5% nucleotide identity in ORF1 and 99.8% in ORF2, suggesting that the sequences belong to the same virus species.

In the ORF1 coding region, nucleotide identities to reference sequences ranged from 74.4% to 82%, with the highest similarity to sequences from Sierra Leone and Spain (PQ541186, PQ537344). In the ORF2 region, identities ranged from 65.5% to 86.3%, with the highest similarity to a sequence from Germany (GU345043). The previously reported Kenyan rat-HEV sequence (PV861667, Kenya 2016) showed 81.1–81.3% identity in ORF1 and 85.2% in ORF2, indicating that our *Rattus norvegicus*–derived sequences are related to, but genetically distinct from this earlier Kenyan strain. Phylogenetic analyses based on ORF1 and ORF2 nucleotide sequences showed that the sequences obtained in this study clustered within the *Rocahepevirus ratti* clade. In both trees, the sequences formed a strongly supported sub-cluster within a broader assemblage of rat-HEV strains from geographically diverse regions, consistent with the limited geographic structuring reported for rat-HEV (Fig. 2).

**Figure 2.**
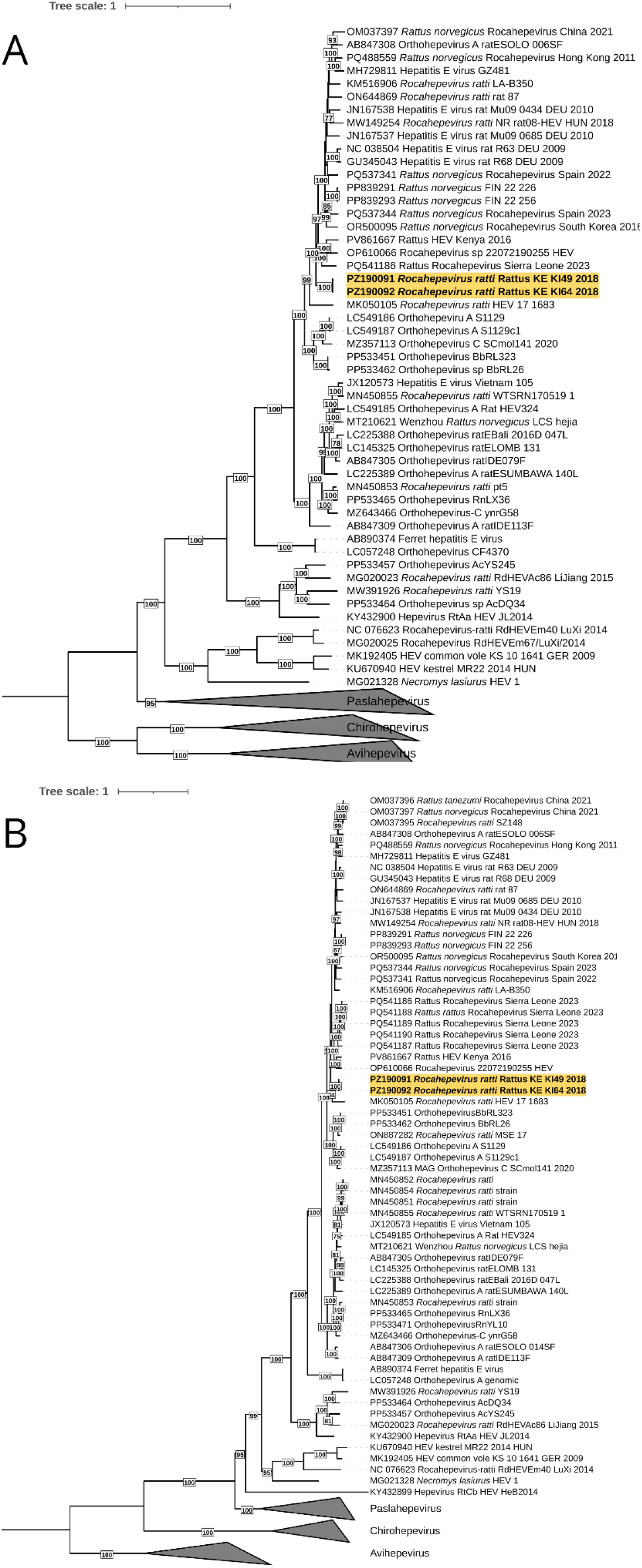
A maximum-likelihood phylogenetic tree of ORF1 (A) and ORF2 (B) nucleotide sequences of hepeviruses. The trees are midpoint-rooted, bootstrap values ≥75 are shown, and the sequences from this study are highlighted in yellow. The best-fit substitution models (GTR+F+R7 and TIM2+F+I+R5, respectively) were selected by ModelFinder [36] using the BIC, and trees were inferred in IQ-TREE [37] with 1,000 ultrafast bootstrap replicates. Visualization was conducted using iTOL [38].

Comparative genome analysis revealed sequence length variation within ORF1. Multiple sequence alignment with reference *Rocahepevirus ratti* showed that, although small insertions and deletions were observed across multiple rat-HEV sequences, the two Kenyan genomes (KI49 and KI64) shared distinct, larger deletions within the same region. Both genomes contained a 93-nt in-frame deletion, while an additional 15-nt in-frame deletion was present in KI64. In addition to these larger deletions, smaller deletions of 2 nt (in KI49) and 17 nt (in KI64), relative to the reference sequence, resulted in frameshifts within ORF1. The positions of these deletions are shown in Supplementary Figure 5. The deletion junctions were confirmed by PCR amplification followed by Sanger sequencing (Supplementary Material S3), supporting the authenticity of these variations. More detailed description of NGS data and genome analysis is described in Supplementary Materials (Supplementary Table 1, Supplementary Figures 1-5).

### Multiple paramyxoviruses detected across diverse host species

In total nine animals, *Acomys ignitus* (A80), *Crocidura sp* (A50 and A53), *Mus minutodes* (A49, A52 and A56) and *Rattus norvegicus* (KI147, KI170, KI171), tested positive for paramyxoviruses in kidney tissue with the pan-jeilong RT-qPCR. Three of the positive animals were also positive in lung tissue (A49, A52, A80). Sanger sequences were obtained from 8 individuals, while one (A53) remained negative, likely due to lower viral load indicated by a higher Ct-value (38.7) compared to the other samples (Ct-values 23.0-29.0). One positive sample (A80) was selected for NGS with metagenomic approach with Illumina, as the kidney sample had a relatively low Ct-value (29.4) and both kidney and lung tissues were positive.

Three of the sequences were acquired from *Rattus norvegicus* individuals, from which KI170 and KI171 shared high similarities with known Beilong virus strains (90.91-96.97% nt-identity), while KI147 had 97.30% nt-identity with a *Rattus tanezumi* jeilongvirus from Singapore. In the maximum-likelihood phylogenetic tree (Fig 3), KI170 and KI171 sequences are placed in the same branch with a previously detected Beilong virus strain, and KI147 is placed in the same branch with jeilongviruses from *Rattus tanezumi* and *Bandicota indica*.

**Figure 3.**
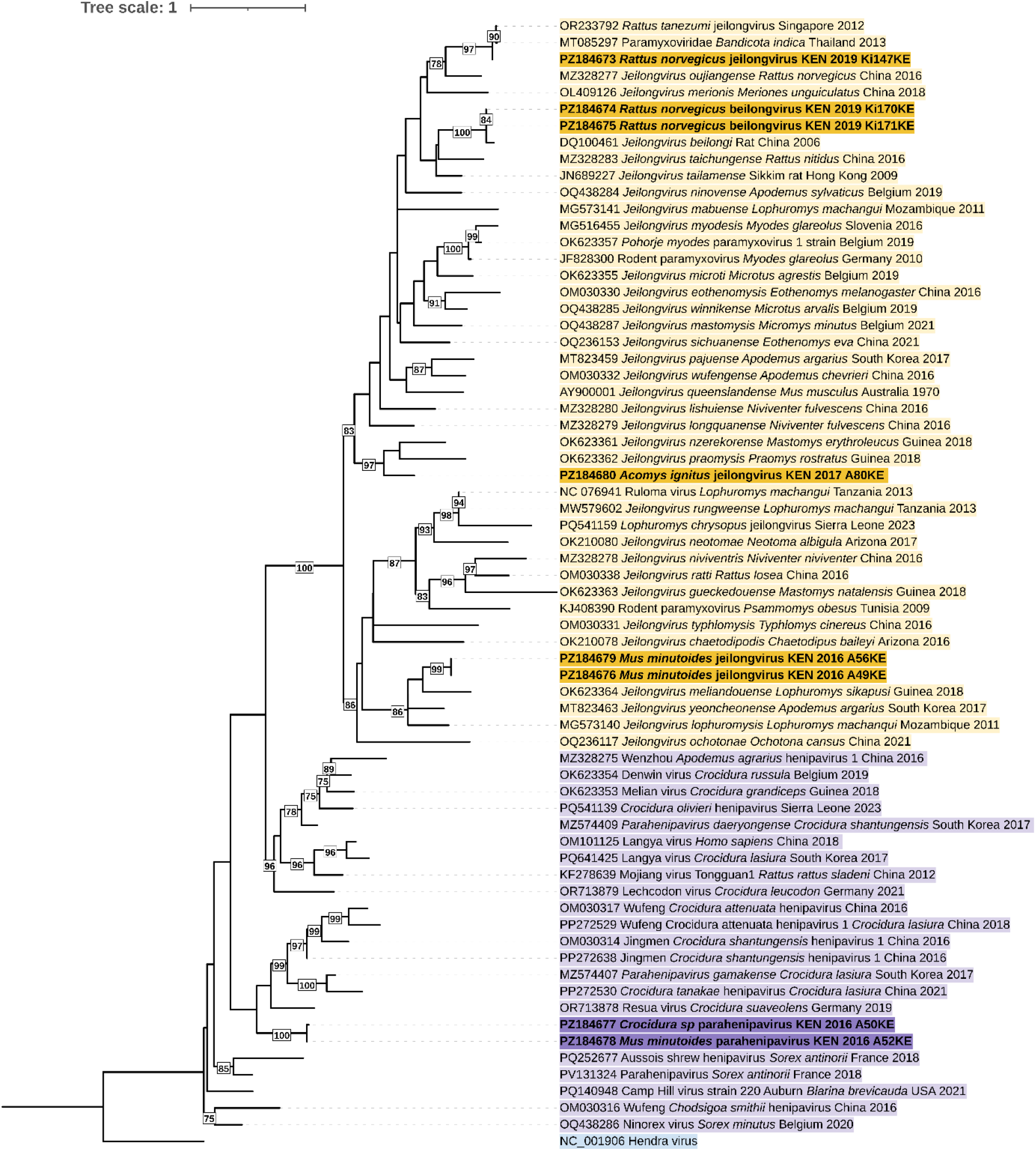
The phylogenetic tree of 234-nt long paramyxovirus sequences rooted with Hendra virus. Jeilongviruses and parahenipaviruses are indicated with yellow and purple color, respectively. Sequences from this study are bolded and colored with darker color. The maximum-likelihood tree is constructed using the GTR+F+I+G4 fitting model according to BIC with 1000 bootstrap replicates, using the IQTREE ModelFinder[36] and IQTREE [37]. Bootstrap values ≥75 are shown. Visualization of the was conducted using iTOL [38]

Five other sequences had less nucleotide identity with known paramyxoviruses available in the NCBI GenBank. Sequences A50 from *Crocidura sp* and A52 from *Mus minutoides* were highly similar to each other with 98.72% nt-identity. BLAST nucleotide searches indicated that both shared highest identity with parahenipaviruses, closest being the Jingmen Crocidura shantungensis henipavirus (PP272638.1) with 76.47% and 77.59% nt-identity, respectively. In the phylogenetic tree, these sequences are placed in a separate highly supported branch within the parahenipavirus clade.

The two sequences from *Mus minutoides* (A49 and A56) were identical. They clustered within the jeilongvirus clade and shared 76% nt-identity with the Ninapo virus (*Jeilongvirus ninovense*, OQ438284) from Belgium. The sequence from *Acomys ignitus* (A80) shared 76.47% and 74.77% nt-identity with Lophuromys chrysopus jeilongvirus (PQ541159.1) and Ruloma virus (NC_076941.1), respectively. Compared to the currently classified jeilongvirus species recognized by ICTV, it shared 72.07% nt-identity with *Jeilongvirus nzerekorense* (OK623361.1) and 65.77% *Jeilongvirus praomysis* (OK623362.1). In the phylogenetic tree, the sequence forms a distinct branch with *Jeilongvirus praomysis* and *Jeilongvirus nzerekorense* (Fig 3).

Sample A80 was selected for metatranscriptomic nextgeneration sequencing, which yielded a nearly complete jeilongvirus genome, here tentatively named as *Acomys ignitus* jeilongvirus. This sample was selected based on its divergence, low Ct-value and phylogenetic position in 234-nt tree (Fig 3). Remaining gaps were subsequently closed using targeted conventional PCRs followed by Sanger sequencing (Supplementary table 2). This combined approach resulted in a complete 17,332-nt coding sequence. Based on the BLAST annotation, the closest viral sequence was Meliandou praomys virus (*Jeilongvirus praomysis*, OK623362) with 76.64% nt-identity (Fig 4). The genome organization of *Acomys ignitus* jeilongvirus follows the typical jeilongvirus gene order and encodes nucleocapsid protein (N), P/V/C proteins, matrix protein (M), fusion protein (F), transmembrane protein (TM), attachment glycoprotein (G) and RNA polymerase (L) (Fig 5). The maximum-likelihood phylogenetic trees of each protein are shown in Supplementary Figures 6-12. Based on the similarity plot analysis (Fig 5), A80 shows major dissimilarities in the gene areas of N, TM and L proteins compared to closest jeilongviruses.

**Figure 4.**
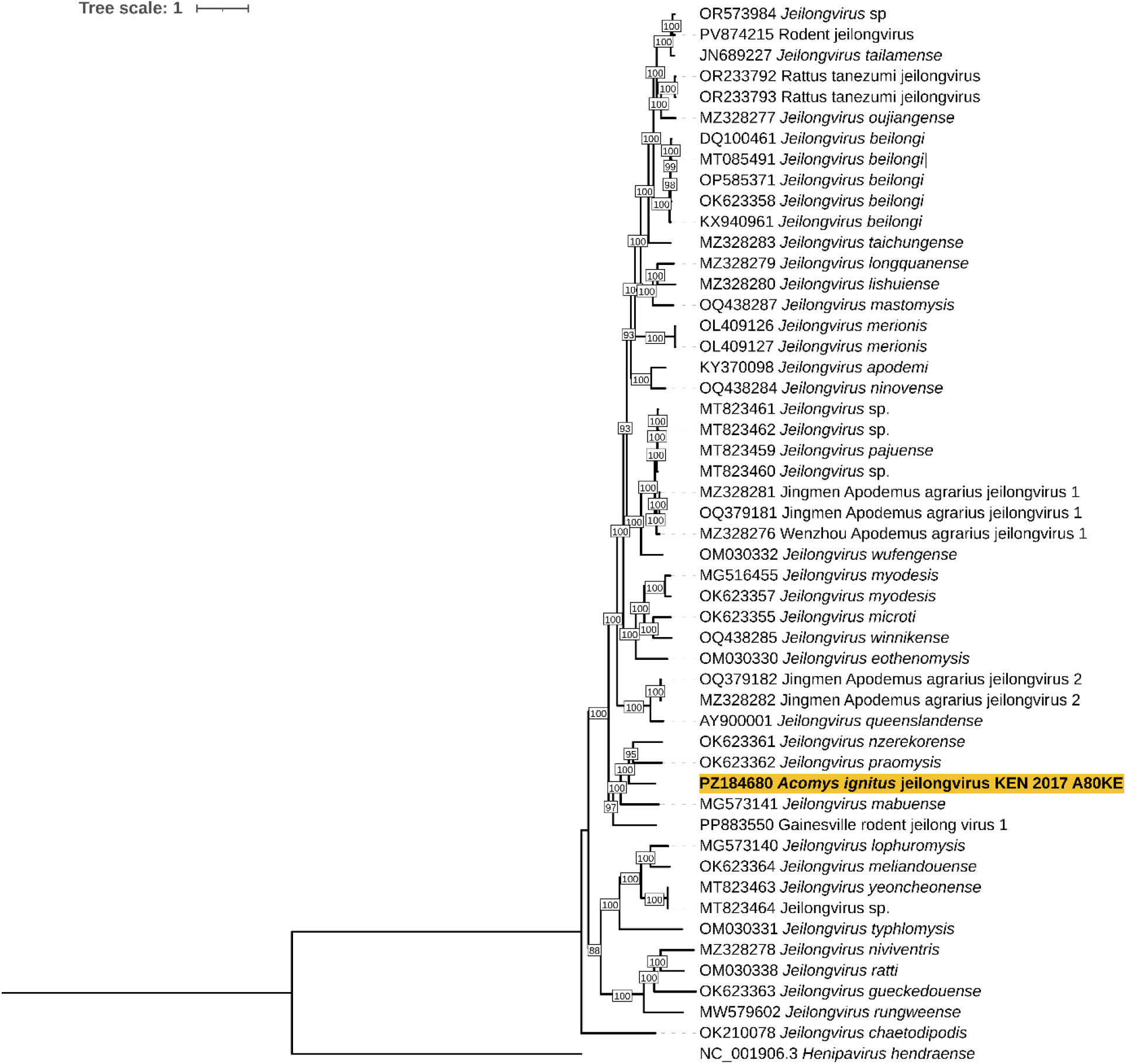
A maximum-likelihood phylogenetic tree of A80 *Acomys ignitus* jeilongvirus with other full-genome jeilongvirus sequneces from NCBI databank. The tree is constructed using GTR+F+R8 model with IQTREE with 1000 bootstrap replicates and rooted with Hendra virus. Visualization of the was conducted using iTOL [38].

**Figure 5.**
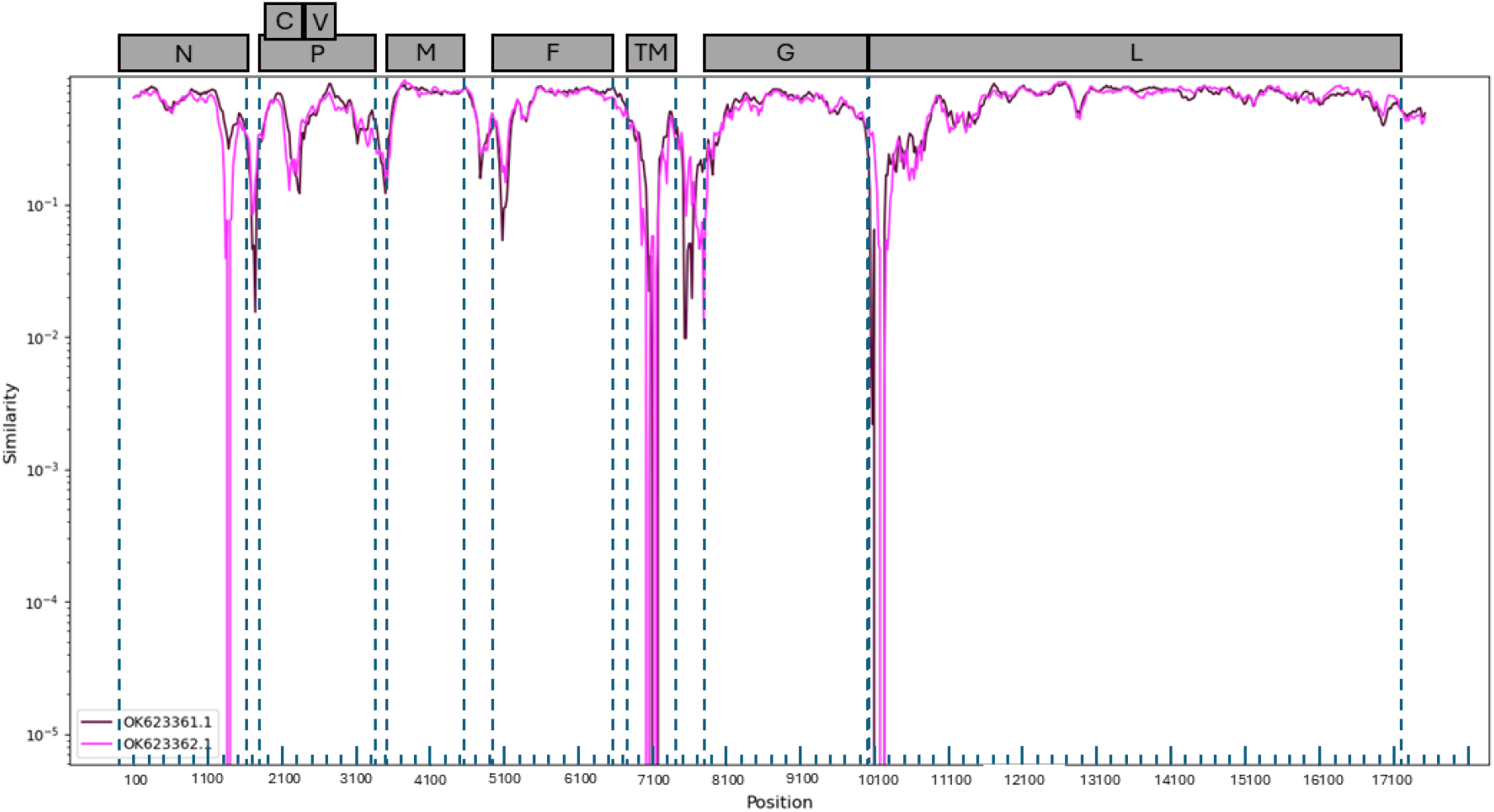
A similarity plot analysis of the closest currently classified jeilongvirus species, *Jeilongvirus nzerekorence* (NCBI: OK623361) and *Jeilongvirus praomysis* (NCBI: OK623362), against *Acomys ignitus* jeilongvirus. SimPlot was constructed using SimPlot++ [39] with 200 bp sliding window and 20 bp steps with Jukes-Cantor substitution model. The ORFs of A80 are indicated in the top part with grey boxes and marked with dashed lines in the plot.

## Discussion

Characterized by high population densities and expansive ecological niches, rodents and shrews represent critical interfaces for zoonotic spillover. While the high viral richness within Rodentia is well-documented[5], the recent identification of novel pathogens in Soricomorpha (shrews) [40], underscores their emerging importance as vital targets for systematic biosurveillance. This study contributes to addressing the lack of pathogen screening studies comparing rural and urban settings in Kenya, despite earlier research demonstrating the presence of both known and novel viruses in the country [14–17]. In this study, we identified rat-HEV in the Kibera informal settlement, while paramyxoviruses were detected in the rodents and shrews from both the Kibera informal urban settlement and the Taita rural region.

HEV RNA was detected in both *Rattus norvegicus* and *Mus musculus* from the Kibera informal settlement. The sequences clustered within the species *Rocahepevirus ratti* (genus *Rocahepevirus*) and belonged to the HEV-C1 lineage (formerly classified as *Orthohepevirus C genotype 1*), which is distinct from Paslahepevirus balayani genotypes typically associated with human infections [41,42]. Rat-HEV is becoming a zoonotic pathogen of growing public health concern, and the confirmed human infections resemble classic acute hepatitis. As of January 2026, at least 48 human infections have been reported [43]. Its transmission route remains poorly understood, but it is suspected to be transmitted through contaminated environment, food or water, thus making it a virus of interest for further studies[41].

Globally, literature on rat-HEV detection in *Mus musculus* is very limited. One documented instance has been reported in *Mus musculus* from China [44]. The few detections of rat-HEV in *Mus sp*, compared with its frequent detection in *Rattus spp*.[45], suggest that infections in *Mus musculus* most likely represent spillover cases. Also, the fact that the NGS failed suggests a lower viral load typical for spillover cases. In the phylogenetic analyses, the sequence obtained in this study clustered among rat-HEV strains from multiple geographic regions, consistent with the lack of clear geographic structuring observed for rat HEV.

The detection of the rat-HEV in Kibera informal settlement reflects the ubiquity of rodent reservoirs in densely populated urban areas. The detection of the virus in different localities within Kibera raises concern about the possibility of the virus spilling over to humans [20]. Rodents may also play a critical role in the spread of the virus to humans as they forage in houses; however, due to limited diagnostic capacity within the country, such pathogens may remain undetected or unrecognized in human population.

Paramyxoviruses have previously been reported in small mammals from the families *Muridae, Cricetidae, Sciuridae* and *Soricidae* [26,46]. In line with these observations, we detected paramyxoviruses in multiple rodent and shrew species across both study sites, suggesting that these viruses are circulating in diverse small mammal communities in Kenya. Similar patterns have been described in earlier studies carried out within the country [16,17]. Paramyxoviruses are generally considered to show a degree of host association and are mainly transmitted through direct contact or airborne routes [47]. The family includes more than 70 recognized species, some of which are known to infect humans [48].

The paramyxoviruses detected in this study cluster into two paramyxovirus genera: *Jeilongvirus* and *Parahenipavirus*. The sequences from *Rattus norvegicus* (KI147, KI170, KI171) form sub-clusters with other rat-related viruses within the genus *Jeilongvirus*. This pattern is consistent with the previously observed tendency of jeilongviruses to show host-associated clustering.

Two sequences obtained from *Mus minutoides* (A49 and A56) formed a distinct and divergent cluster in the short-fragment phylogenetic analysis, suggestive of a putative novel virus species; however, these sequences were not subjected to further sequencing. In contrast, the sequence from *Acomys ignitus* (A80) was confirmed as a novel virus species based on complete coding sequence analysis. Sequences from *Crocidura sp* (A50) and *Mus minutoides* (A52) cluster together in the genus *Parahenipavirus*, which includes the zoonotic Langya virus [7]. However, these short Sanger sequence results need to be confirmed with NGS. These two highly similar sequences between a shrew and mouse trapped from the same site support the previous finding that jeilongviruses can transmit between host orders [49].

The presence of viruses belonging to two different paramyxovirus genera supports the view that rodents and shrews harbor a diverse range of paramyxoviruses. A complete coding sequence was obtained from *Acomys ignitus* (A80), and this sequence was genetically distinct from known jeilongvirus species with nucleotide identity of 65.77-76.47%. Based on the results of the phylogenetic tree (Fig 4) and the similarity plot analyses (Fig 5) *Acomys ignitus* jeilongvirus likely represents a novel jeilongvirus species. The genome organization of *Acomys ignitus* jeilongvirus follows that of the closest jeilongvirus species [26], *Jeilongvirus nzerekorense* and *Jeilongvirus praomysis*, but has major differences in the nucleotide sequence of three different genes. Separate phylogenetic trees constructed with amino acid sequences of different jeilongvirus proteins (Supp. Figure 1-7) consistently placed *Acomys ignitus* jeilongvirus on a distinct branch with *Jeilongvirus nzerekorense, Jeilongvirus praomysis* and *Jeilongvirus mabuense* as its closest relatives. These viruses have been detected in Guinea and Mozambique from different rodent species [26]. The closest relative to *Acomys ignitus* jeilongvirus sequence is the *Jeilongvirus nzerekorense*, which was predicted to possess high zoonotic potential based on its genomic features using machine-learning method [50], highlighting the need for further investigation of these viruses. However, no human infections of jeilongviruses have been reported to date.

According to the ICTV classification criteria, phylogenetic analyses based on the amino acid sequence of the L gene are used for species demarcation [47]. In this context, *Acomys ignitus* jeilongvirus forms a distinct lineage with *Jeilongvirus praomysis* separate from other jeilongviruses supported by a bootstrap value of 77 (Supplementary Fig. 7), providing evidence for their common ancestor.

In conclusion, Kenyan rodents and shrews host a diversity of viruses. We detected three rat-HEV and nine paramyxoviruses, highlighting the role of small mammals as important reservoirs. The detection of rat-HEV in Kibera rodent populations underscores the potential for zoonotic spillover in densely populated informal urban settlements. The discovery of a novel *Acomys ignitus* jeilongvirus and other divergent paramyxoviruses from both urban and rural sites further demonstrates the complexity of viral circulation in wildlife. To date no human infections of the detected viruses have been reported in Kenya, highlight the need for continued One Health surveillance and expanded genomic sequencing studies to better understand the zoonotic risk and strengthen public health preparedness in the country experiencing rapid land-use change and subsequent increased exposure to rodent-borne pathogens.

## Abbreviations

HEV: Hepatitis E Virus
ORF: Open reading frame
ICTV: International Committee on Taxonomy of Viruses

## Acknowledgments

We acknowledge field assistants at the Taita research station for their support in rodent trapping and assisting in shipment of the samples from Taita to Nairobi. We acknowledge the Taita and Kibera local community for allowing us to trap samples within their premises. We acknowledge the KAVI lab technicians for their enormous support in helping to sort and pack the samples for shipping from Nairobi to University of Helsinki, Finland. We acknowledge Simo Miettinen and Lauri Kirjalainen (University of Helsinki) for their support during the lab work, especially the sample retrieval and curation. We acknowledge Leena Palmunen (University of Helsinki) for the guidance and help in the NGS library preparation.

## Permits and funding

This research was funded by Jenny and Antti Wihuri Foundation, Research Council of Finland (grant no. 358323 and 339510 to Sironen, grant no. 318725 to Vapalahti), the European Union ZOOSURSY (CAN700002203) project, HUS Diagnostic Center (Helsinki University Hospital, TYH2025225), the Swedish Research Council (grant number 2022-03218) and Sigrid Jusélius Foundation (to Sironen and Vapalahti). This research was also partly supported by funding from the European Union’s EU4Health programme under Grant Agreement Nr 101102733 DURABLE. Views and opinions expressed do not necessarily reflect those of the European Union or HaDEA. Neither the European Union nor the granting authority can be held responsible for them.

## Author contributions

- Conceptualization: Sironen T, Korhonen E, Omu A, Olli V
- Data curation: Pulkkinen E, Altan E, Ochola G
- Formal analysis: Ochola G, Pulkkinen E, Altan E
- Funding acquisition: Sironen T, Jääskeläinen AJ, Lindahl J, Forbes K
- Investigation: Sironen T, Forbes K, Laakkonen J, Ogola J, Mweu A, Uusitalo J, Masika M
- Methodology: Sironen T, Korhonen E, Pulkkinen E, Ochola G, Altan E, Jääskeläinen AJ, Forbes K, Lindahl J, Vauhkonen H, Ogola J, Masika M, Omu Anzala, Smura T,
- Project administration: Sironen T, Forbes K, Omu A, Olli V
- Resources: Sironen T, Jääskeläinen AJ
- Supervision: Sironen T, Korhonen E, Altan E, Jääskeläinen AJ
- Validation: Sironen T, Jääskeläinen AJ, Smura T
- Visualization: Pulkkinen E, Mäkelä H, Ochola G
- Writing – original draft: Ochola G, Pulkkinen E, Altan E, Sironen T
- Writing – review and editing: All the authors

