## Supplementary material for "Rat hepatitis E virus and novel paramyxoviruses in synanthropic rodents and shrews in Kenya": Supplimentary materials

**Supplementary Materials**

**S1. Detailed Description of Study Sites and Spatial Data Processing**

**Study site characterization**

Two ecologically distinct sites in Kenya were included in this study: Kibera informal settlement (Nairobi County) and the Taita Hills region (Taita-Taveta County).

Kibera is located approximately 5 km southwest of Nairobi city center. It is characterized by high human population density, informal housing structures, limited sanitation infrastructure, inadequate waste management systems, and the absence of formal sewerage networks. These environmental conditions create favorable habitats for synanthropic rodents and increase opportunities for human–rodent interactions.

The Taita Hills region forms part of the Eastern Arc Mountains in southeastern Kenya, a recognized biodiversity hotspot. Historically covered by continuous montane forest, the landscape is now highly fragmented due to agricultural expansion and human settlement. The region consists of indigenous forest patches, mixed forests, and plantation areas embedded within densely populated agricultural landscapes. The proximity of forest fragments to farms and rural settlements may facilitate wildlife–livestock–human contact.

**Spatial data processing and map generation**

Maps illustrating the sampling locations were generated using QGIS software (version 3.36.2).

Base layers including country borders, administrative boundaries, water bodies, roads, railways, and buildings were obtained from Natural Earth (2025 release) and OpenStreetMap (2025 data extract). True-color background imagery for the Taita Hills region was derived from Sentinel-2 MSI Level-1C satellite imagery (ESA/Copernicus; tile T37MCR; acquisition date: 09 September 2025).

Sampling locations were georeferenced using handheld GPS devices during field collection. Positive detection sites were visualized using distinct color coding to differentiate overall sampling points and virus-positive locations.

**Rationale for site selection**

The two study areas were selected to represent contrasting ecological and socio-environmental settings:

- An urban informal settlement with high human density and intense rodent–human contact (Kibera)
- A fragmented forest–agricultural interface characterized by biodiversity and land-use change (Taita Hills)

This design enabled comparison of viral detection across urban and rural ecological gradients within a One Health framework.

**S2. NGS library preparation**

The sequencing libraries were prepared using NEBNext® Ultra™ II RNA Library Prep Kit for Illumina® (New England Biolabs, Ipswich, MA, USA) system with slight modifications. The ribosomal RNAs were depleted from approximately 500 ng of nanodrop-quantified TRIzol-extracted total RNA using NEBNext® rRNA Depletion Kit (Human/Mouse/Rat, New England Biolabs) with half of the recommended reaction volumes. The rRNA-depleted material was purified using RNAClean XP magnetic beads (Beckman Coulter, Indianapolis, IN, USA). The first and second strand reactions were carried out as recommended and five minutes was used for the fragmentation time. SPRIselect magnetic beads (Beckman Coulter) and 12 uL elution volume was used in the subsequent purification of the double-stranded cDNA. The End Prep and adapter ligation steps were carried out in 1/5 of the recommended volumes using a 1/5 dilution of the adapter. After SPRIselect bead purification the adapter-ligated cDNA was eluted in 15 uL of sterile water and 11.5 uL was used in the final amplification reaction using 12.5 uL of Q5 Master mix (New England Biolabs) and 1 uL of CleanPlex Plated Unique Dual-Indexed PCR Primers for Illumina (Paragon Genomics, Fremont, CA, USA) and 13 cycles of PCR amplification. fo. The PCR products were purified using SPRIselect beads and eluted in 20 uL of sterile water. Library sizes were approximated on agarose gel, concentrations determined with Qubit dsDNA High sensitivity system (Thermo Fisher Scientific, Waltham, MA, USA) and run on Illumina NovaSeq X sequencer (Illumina, San Diego, CA, USA) at the Institute for Molecular Medicine Finland FIMM Genomics unit using 25B 300c sequencing kit.

**Supplementary Table 1**. NGS results

| Sample ID | Total Reads | After filter total read | Q30 bases | Virus read % | Jeilongvirus | Rocahepevirus ratti | Marmot Picobirnavirus |
| --- | --- | --- | --- | --- | --- | --- | --- |
| KI_49 | 237.63.110 | 237.633.080 | 98.4 | 0.07 |  | 154 |  |
| KI_64 | 78.000.000 | 67.500.000 | 89.2 | 0.2 |  | 3954 |  |
| A80 | 326.509900 | 326509488 | 98.2 | 0.04 | 1493 |  | 6 |

**Supplementary Table 2.** Table of primers designed and the PCR conditions to fill the missing gaps in the NGS data (PMV-Paramyxovirus, HEV- Hepatitis E virus

| virus | primer_Name | Sequence 5’-3’ | Sequence L | Size | Tm |
| --- | --- | --- | --- | --- | --- |
| PMV | LUA80_first_4,349 F | CCTTCATGTGACCGTTGCAG | 20 | 795bp | 58 |
|  | LUA80_first_5,144 R | GCTGCTCCTGATATTTTCAACCC | 23 |  |  |
|  | LUA80_first_7,371 F | GGCTATTCCAACAATAATACCACC | 24 | 528bp | 56 |
|  | LUA80_first_7,899 R | GCAAGCATAGTTTTGCATGTTG | 22 |  |  |
|  | LUA80_first_10,295 F | CCTTTGGCTACAGCACCTGA | 20 | 517bp | 56 |
|  | LUA80_first_10,812 R | ACTTACCTGTAACTATGGGGC | 21 |  |  |
| HEV | HEV 2276F | GATTACAGACGGCGGAGAGA | 20 | 552bp | 57 |
|  | HEV2919R | CTCTTCCCCTTGGAGTGCTG | 20 |  |  |
|  | HEV2290F | TTACAGACGGCGGAGAGATC | 20 | 466bp | 57 |
|  | HEV2854R | GTCGGCACAATGACCAGATC | 21 |  |  |

PCR for filling the missing gaps was performed in a 25 µL reaction containing 1× AllTaq™ PCR Master Mix (QIAGEN, Hilden, Germany), 0.4 µM each of forward primer and reverse primer, 6.25 µL master mix, 1 µL of cDNA template, and top up with nuclear free water. Thermal cycling conditions were an initial denaturation at 95 °C for 2 min, followed by 40 cycles of 95 °C for 15 s, annealing at (3 °C below the lower primer TM) for 30 s, and extension at 72 °C for 30 s, with a final extension at 72 °C for 10 min, infinite hold at 4 °C.

The Initial conventional PCR screening for poxvirus, hantavirus and coronavirus was done using Phusion Flash High‑Fidelity PCR Master Mix (Thermo Fisher Scientific, Waltham, MA, USA), 0.4 µM each primer, 15 µL master mix, 1 µL of cDNA template, and top up with nuclear free water. Thermal cycling conditions were an initial denaturation at 98 °C for 1 min, followed by 35 cycles of 98 °C for 5 s, annealing at (lower primer TM+3 °C) for 10 s, and extension at 72 °C for 15 s, with a final extension at 72 °C for 5 min, infinite hold at 4 °C.

**Supplementary Table 3.** NCBI GenBank accession codes of the detected virus sequences. HEV = rat hepatitis virus, PMV = paramyxovirus. SRA data accession code

| HEV | | PMV | |
| --- | --- | --- | --- |
| Sample ID | Accession code | Sample ID | Accession code |
| Ki49 | PZ190091 | Ki147 | PZ184673 |
| Ki64 | PZ190092 | Ki170 | PZ184674 |
| Ki137 | PZ190093 | Ki171 | PZ184675 |
|  |  | A49 | PZ184676 |
|  |  | A50 | PZ184677 |
|  |  | A52 | PZ184678 |
|  |  | A56 | PZ184679 |
|  |  | A80 | PZ184680 |
| SRA data | PRJNA1444134 | SRA data | PRJNA1444134 |

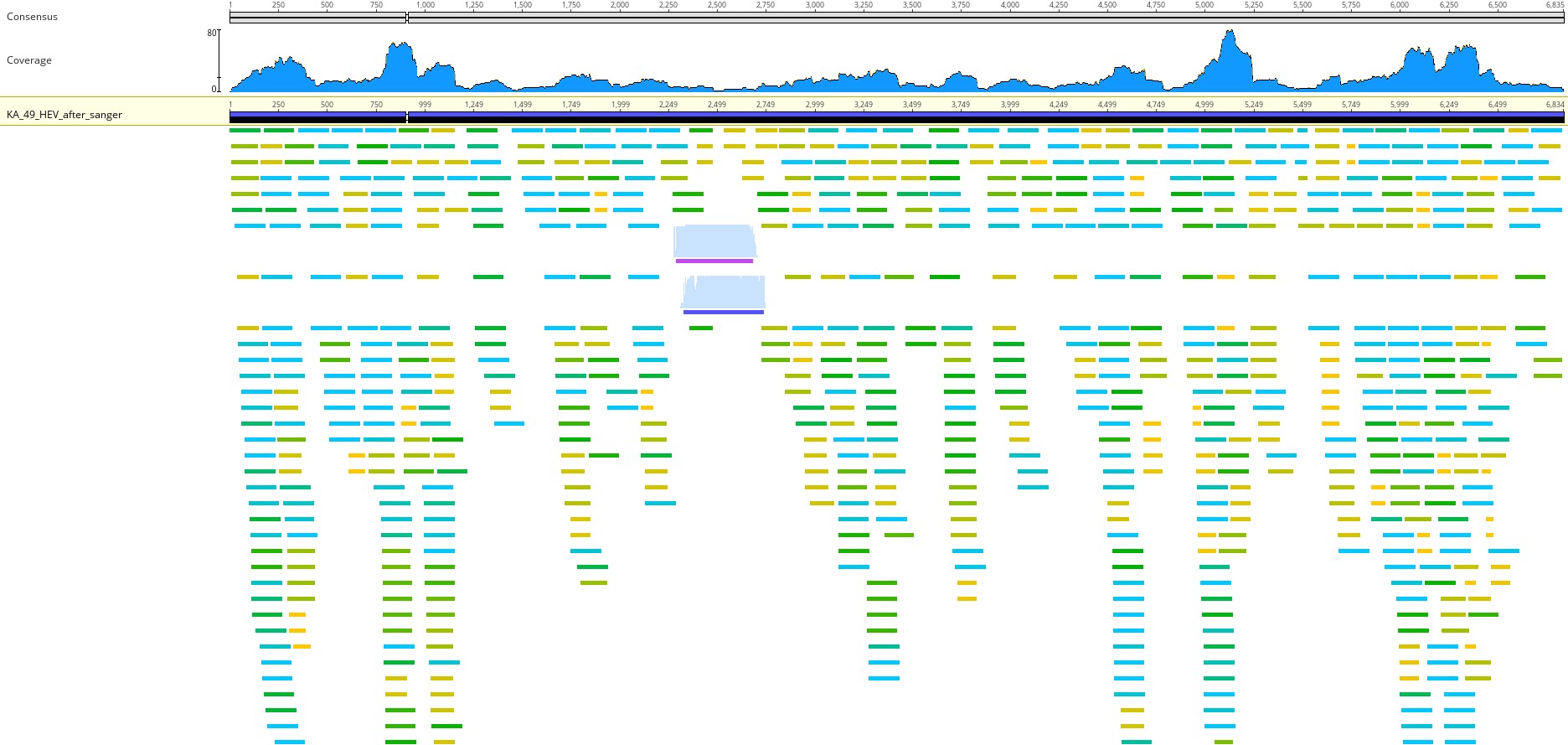

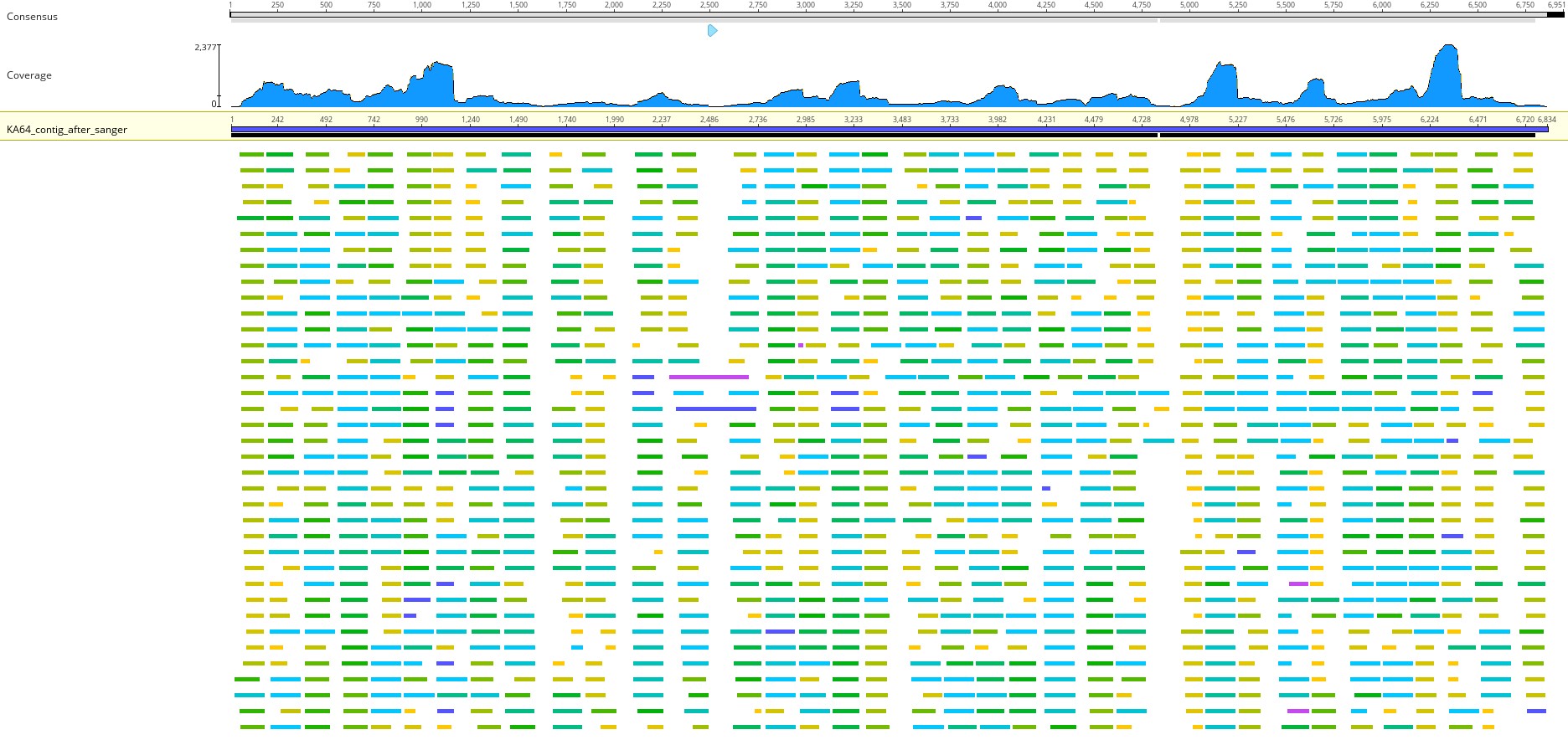
**Supplementary Figure 1**: Genome coverage of rat-HEV

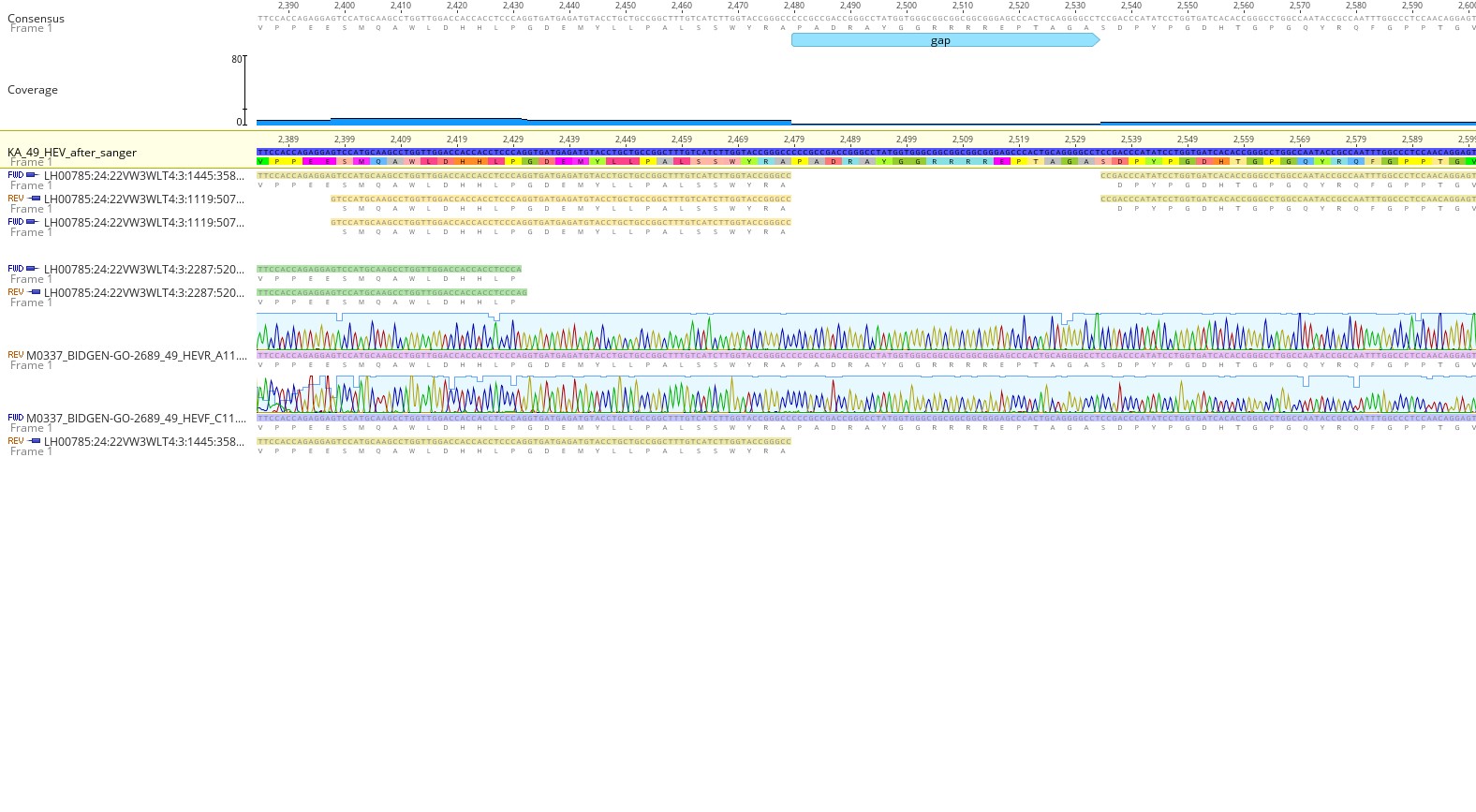
**Supplementary Figure 2**. Confirmation of an NGS-identified genomic gap in the *Rocahepevirus ratti* genome from sample KI49 by Sanger sequencing. The alignment shows the consensus sequence generated from NGS data, read coverage across the region, and the corresponding Sanger sequencing chromatograms obtained using forward and reverse primers. The highlighted region indicates a gap detected in the NGS assembly, which was subsequently resolved by Sanger sequencing, confirming the nucleotide sequence across this region.

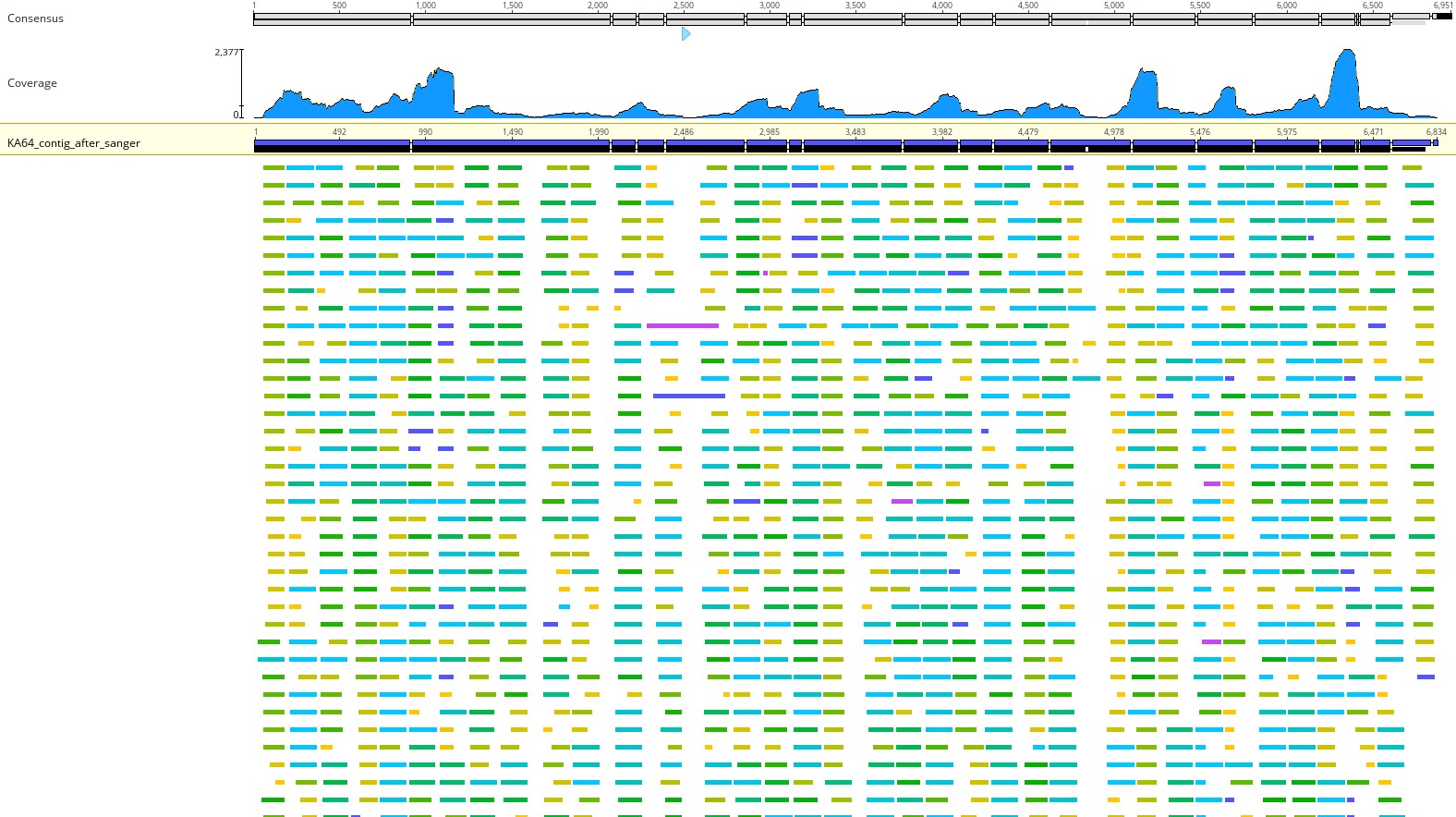
**Supplementary Figure 3**. KI64- NGS Raw read covarages

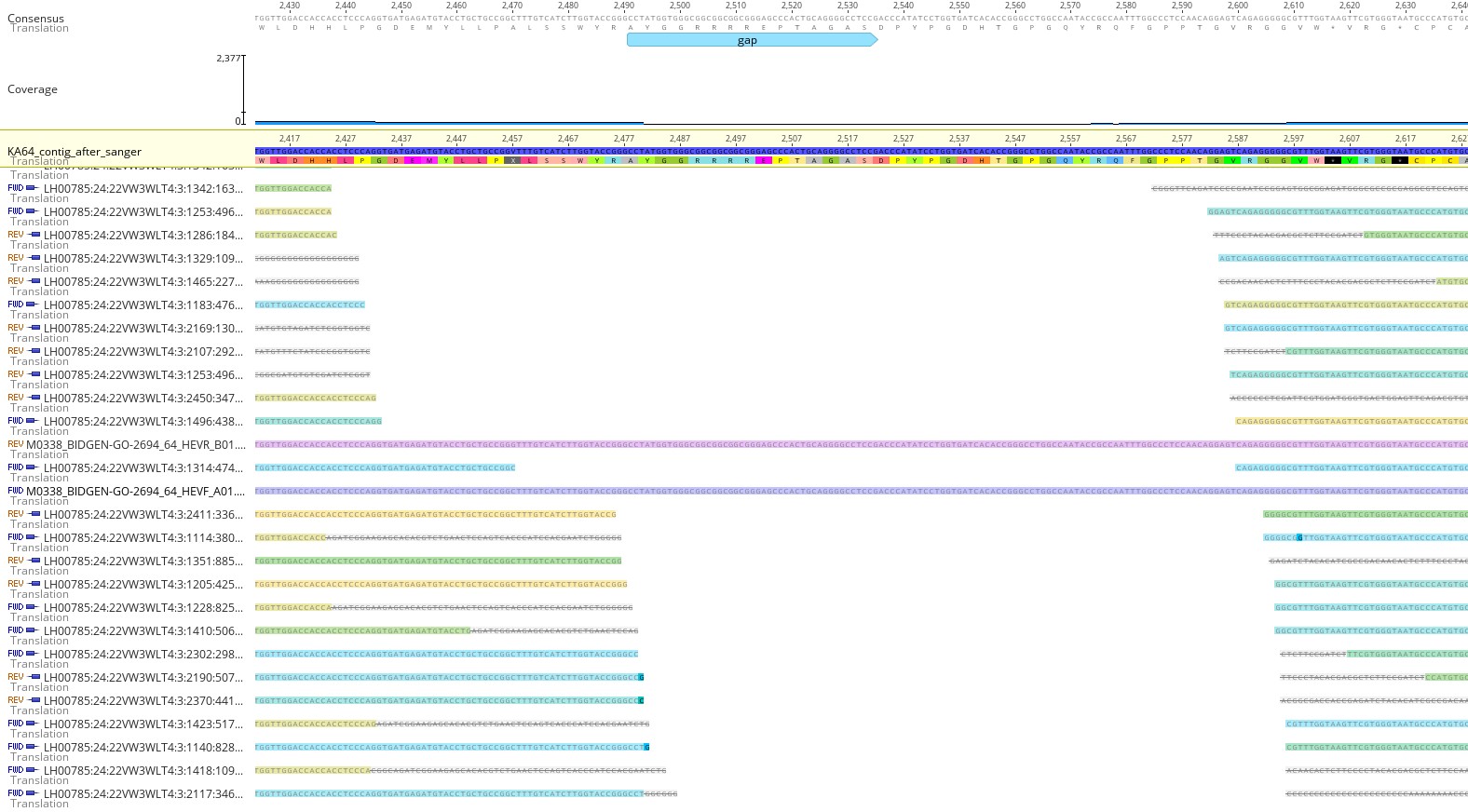
**Supplementary Figure 4.** Mapped Sanger sequence KI64 for filling the gaps

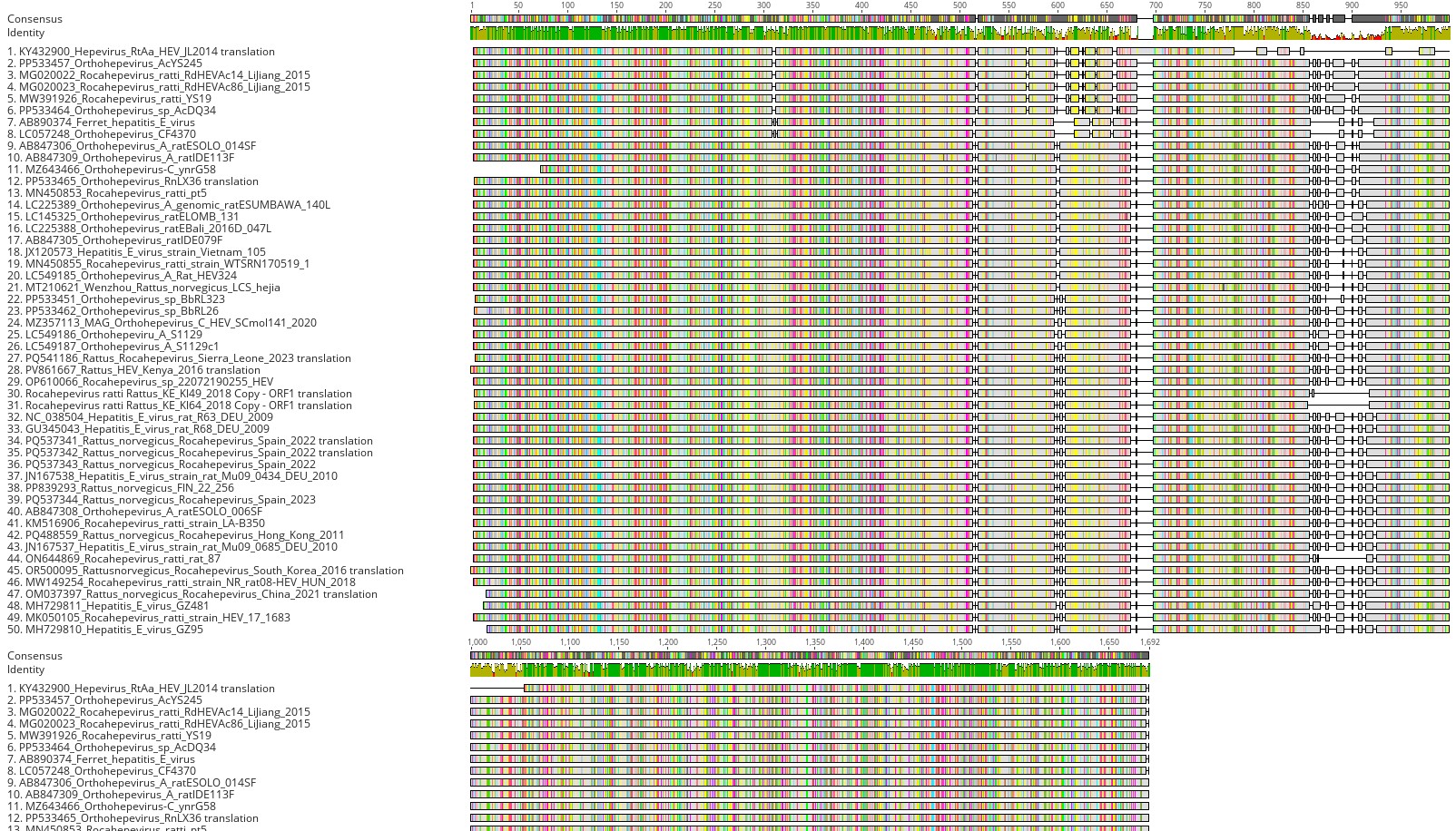

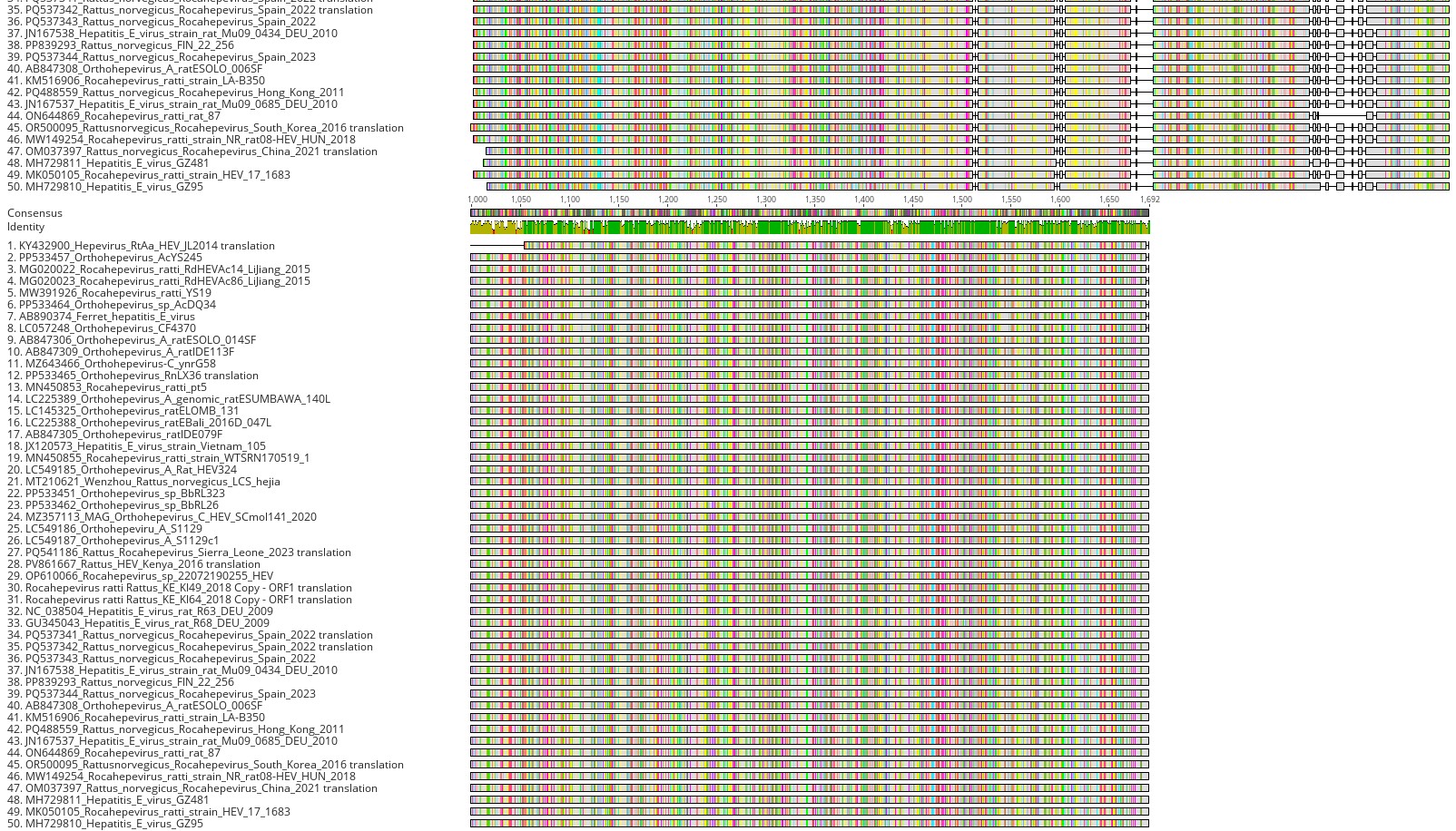
**Supplementary Figure 5** Rocahepevirus ORF1 alignment figure for to show indel rich region

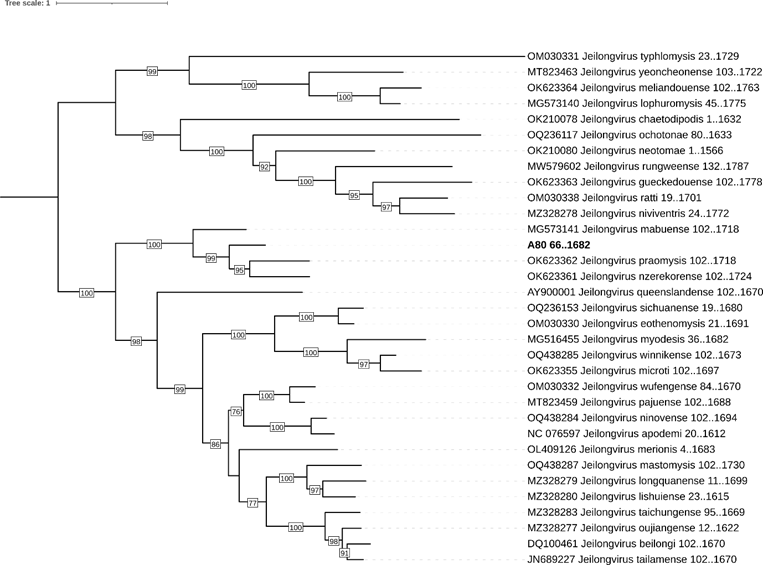

**Supplementary Figure 6** Maximum-likelihood tree of N protein amino acid sequence, constructed using the best fitting model (Q.insect+F+R5) according to BIC with 1000 bootstrap replicates, using the IQTREE ModelFinder[37] and IQTREE[38]. Bootstrap values ≥75 are shown. Visualization of the was conducted using iTOL [39].

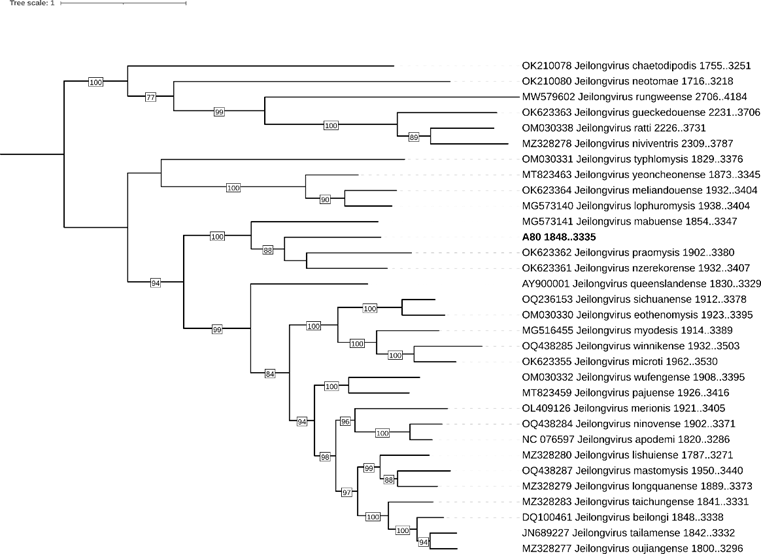

**Supplementary Figure 7** Maximum-likelihood tree of P protein amino acid sequence, constructed using the best fitting model (JTT+F+I+G4) according to BIC with 1000 bootstrap replicates, using the IQTREE ModelFinder[37] and IQTREE[38]. Bootstrap values ≥75 are shown. Visualization of the was conducted using iTOL [39].

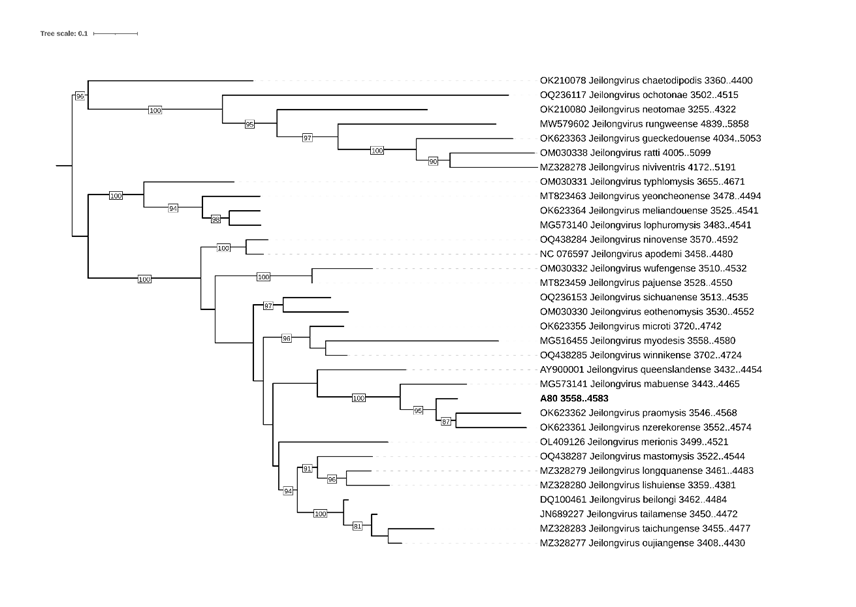

**Supplementary Figure 8** Maximum-likelihood tree of M protein amino acid sequence, constructed using the best fitting model (Q.insect+I+R4) according to BIC with 1000 bootstrap replicates, using the IQTREE ModelFinder[37] and IQTREE[38]. Bootstrap values ≥75 are shown. Visualization of the was conducted using iTOL [39].

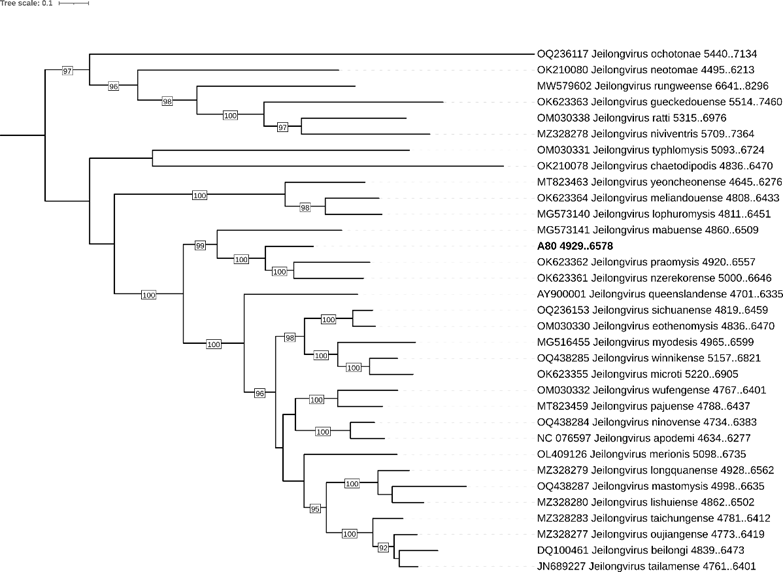

**Supplementary Figure 9** Maximum-likelihood tree of F protein amino acid sequence, constructed using the best fitting model (Q.yeast+I+R5) according to BIC with 1000 bootstrap replicates, using the IQTREE ModelFinder[37] and IQTREEClick or tap here to enter text.. Bootstrap values ≥75 are shown. Visualization of the was conducted using iTOL Click or tap here to enter text..

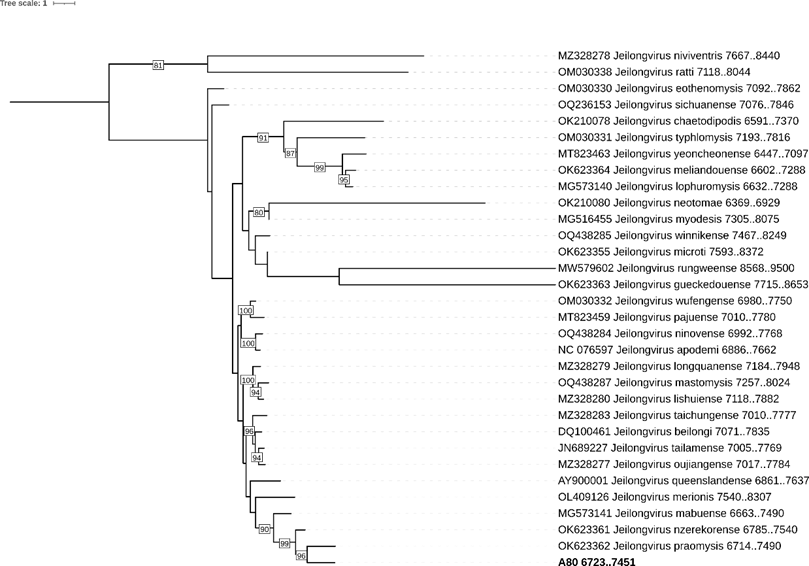

**Supplementary Figure 10** Maximum-likelihood tree of TM protein amino acid sequence, constructed using the best fitting model (Q.insect+F+G4) according to BIC with 1000 bootstrap replicates, using the IQTREE ModelFinder[37] and IQTREE[38]. Bootstrap values ≥75 are shown. Visualization of the was conducted using iTOL [39].

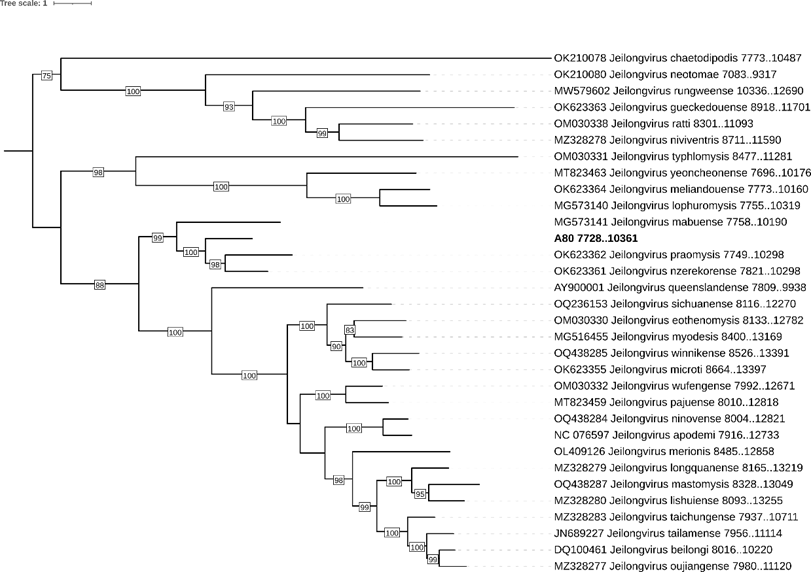

**Supplementary Figure 11** Maximum-likelihood tree of G protein amino acid sequence, constructed using the best fitting model (WAG+F+R6) according to BIC with 1000 bootstrap replicates, using the IQTREE ModelFinder[37] and IQTREE[38]. Bootstrap values ≥75 are shown. Visualization of the was conducted using iTOL [39].

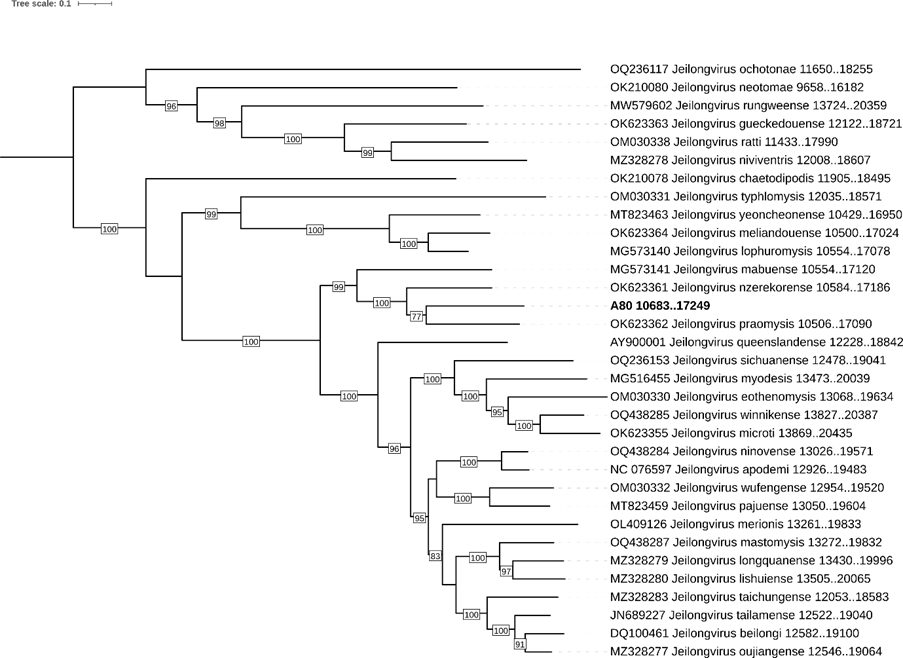

**Supplementary Figure 12.** Maximum-likelihood tree of L protein amino acid sequence, constructed using the best fitting model (Q.yeast+F+I+R5) according to BIC with 1000 bootstrap replicates, using the IQTREE ModelFinder[37] and IQTREE[38]. Bootstrap values ≥75 are shown. Visualization of the was conducted using iTOL [39]
